# Are most human specific proteins encoded by long non-coding RNA ?

**DOI:** 10.1101/2023.11.09.566363

**Authors:** Yves-Henri Sanejouand

## Abstract

By looking for a lack of homologues in a reference database of 27 well-annotated proteomes of primates and 52 well-annotated proteomes of other mammals, 170 putative human-specific proteins were identified. Among them, only 2 are known at the protein level and 23 at the transcript level, according to Uniprot. Though 21 of these 25 proteins are found encoded by an open reading frame of a long non-coding RNA, 60% of them are predicted to be at least 90% globular, with a single structural domain. However, there is a near complete lack of structural knowledge about these proteins, with no tridimensional structure presently available in the Protein Databank and a fair prediction for a single of them in the AlphaFold Protein Structure Database. Moreover, the knowledge about the function of these possibly key proteins remains scarce.

## Introduction

Each time a new genome is sequenced, genes coding for proteins with no known homologue are found, even when genomes of closely related species are available, like in the case of primates [1, 2, 3] or *Drosophila* [4, 5, 6].

In the former case, the hypothesis that human-specific proteins may prove involved in the behavioral or anatomical peculiarities of the human species, like the size of its brain [7] or its functions [8, 9], seems worthy of consideration. So, in the present study, taking advantage of both the high quality of the annotation of the human proteome [10] and of the availability of a significant number of well-annotated primate proteomes [11, 12], an extensive search of human-specific proteins was performed, focusing noteworthy on what can be predicted about their tridimensional structure, with the idea that such a knowledge could provide hints about their function or their origin.

To do so, like in a previous study [13], a reference database was set up. Then, two popular protein structure prediction methods were applied to the putative human-specific proteins thus found, namely, IUPred [14] and AlphaFold [15]. While the former provides a qualitative prediction, stating whether a polypeptide is expected to adopt a globular fold or not, the later is able to predict its tridimensional structure at an atomic level of detail [16]. However, as already underlined previously [13, 15], informations other than the sole sequence of the polypeptide are usually required.

## Methods

### Choice of a reference database

The definition of what a species-specific protein is depends upon what is known about the proteomes of the closest species at a given time. For studying species-specific proteins, as well as for the sake of reproducibility, it is thus important to choose a well-defined reference system [13], that is, a set of reference proteomes. Moreover, since the quality of the annotation of a proteome can vary significantly, both from a proteome to another and as a function of time, these proteomes have to be as well annotated as possible.

To do so, 27 Uniprot reference proteomes [17] of primates were picked, that is, all those that have at least a standard level of annotation^1^, according to the Complete Proteome Detector [18]. Note that 10 of them are high-value outliers, meaning that they have significantly more identified proteins than closely taxonomically related species.

For these 27 reference proteomes, complete BUSCO predictions of single-copy orthologs [19] are found for 94% (median value) of the cases. However, the percentage of short (less than 50 amino-acid residues long) proteins vary widely from a proteome to another, ranging between 0.1% (*Sapejus apella*) and 3.5%, being over 1% in the case of three species only, namely, *Pongo abelii, Pan troglodytes* and *Macaca fascicularis*.

Overall, there are between 19229 (*Chlorocebus sabaeus*) and 50207 (*Macaca fascicularis*) proteins per proteome (40000 ± 7500, on average) in our reference database, with a total of 1.083.746 proteins, including known isoforms.

Figure 1 shows the phylogenetic tree of the primate species considered herein, according to the TimeTree webserver, version 5 [20].

**Figure 1:**
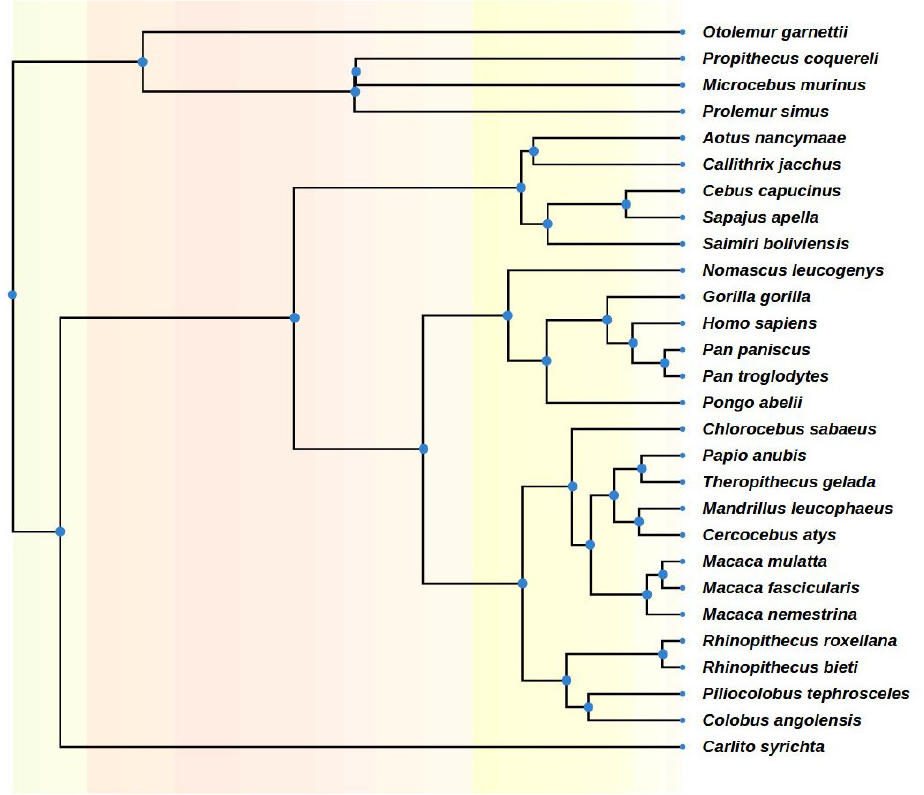
The phylogenetic tree of the primate species with well annotated proteomes considered herein.

### Search of homologues

For each of the 20,449 human proteins at least 30 amino-acid residues long associated to a given gene^2^, homologues in the reference database were looked for using BLAST [21] version 2.6.0+, two proteins being assumed to be homologous when the E-value of their pairwise alignment is lower than 10^−6^ [13, 22, 23]. Note that, in order to avoid an overestimation of the number of specific proteins, due to the filtering of low-entropy segments, that is, of segments of restricted amino-acid composition, composition-based statistics [24] was not considered (-comp based stats 0).

### Non coding RNAs

For each human-specific protein found, that is, for each human protein with no homologue in the reference database, its possible encoding by a human non-coding RNA (ncRNA), as found in the RNAcentral database [25], version 22, was checked, by comparing its sequence to those obtained by translating all at least 90 nucleotides long nonoverlapping open reading frames. Note that it was assumed that both start and stop codons are standard ones, while it is known that peptides encoded by ncRNAs can have atypical stop codons [26].

### Protein globularity

The degree of globularity of each human-specific protein found, that is, the percentage of the protein length predicted to be globular, as well as their number of globular domains, was estimated using the standalone version of IUPred [14, 27]. Note that IUPred performs its predictions by considering the local sequential environment of each aminoacid residue within 2–100 residues in either direction. Note also that, at variance with most recent methods [28], IUPred does not make use of evolutionary data, which are expected to miss in the case of species-specific proteins.

### Structure prediction

Predictions of tridimensional structures were picked in the AlphaFold Protein Structure Database [29]. These structures were obtained with the AlphaFold2 artificial intelligence from DeepMind [15], an algorithm whose predictions proved to be the most accurate of all the submissions for over 90% of the targets of the CASP14 blind prediction experiment [16].

Note that AlphaFold2 can also be used for predicting if a protein has disordered segments [27, 30]. Indeed, AlphaFold2 provides an estimate of the accuracy of its prediction for the position of each amino-acid residue, coined pLDDT^3^, values over 90% corresponding to a high quality and values below 50% to a poor one [29]. Herein, the overall quality of the prediction of the structure of a protein is assumed to be given by the average of the quality of the prediction of the position of its residues ((pLDDT)).

## Results

### How many human-specific proteins ?

The need to use a large enough reference database was assessed as follows. When proteins of *Homo sapiens* not found in the proteome of its closest relative, namely, *Pan troglodytes*, are looked for, 347 of them are identified. Note that this number is lower than a previous estimate obtained ten years ago, namely, 634 [3], maybe as the result of the improvement of the annotation of the human proteome [10]. Indeed, when the search was performed the other way around [13], 1036 chimpanzee-specific proteins^4^ were found.

As a matter of fact, as shown in Figure 2, when the number of proteomes in the reference database is increased, the number of human-specific proteins keeps on dropping, down to 193. Interestingly, a few species have significantly higher contributions to this downsizing, like the fourth (*Pongo abelii*) and tenth (*Macaca fascicularis*), further suggesting that their proteomes are more complete than the others. But while the proteome of *Macaca fascicularis* is indeed the largest in our reference database, the size of the proteome of *Pongo abelii*, with 39491 proteins, is slightly below the average. Indeed, its level of annotation is just considered standard, according to the Complete Proteome Detector [18].

**Figure 2:**
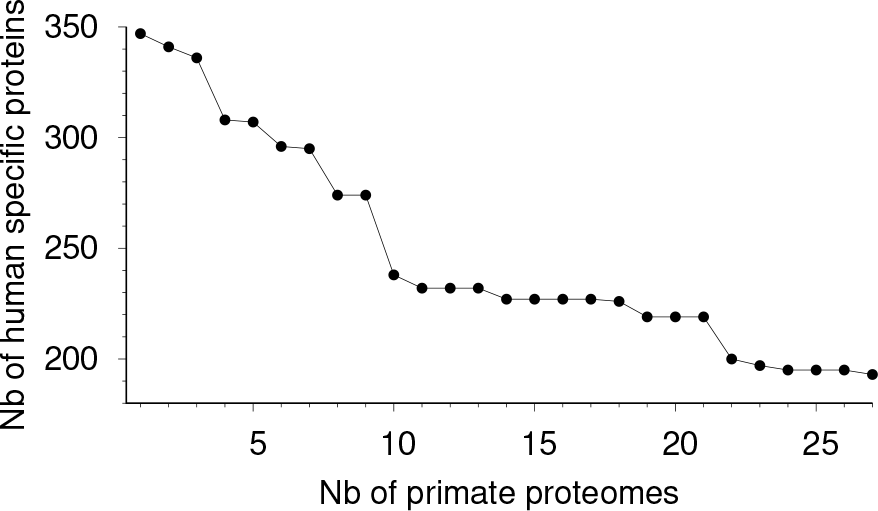
Number of human specific proteins, as a function of the number of primate proteomes in which homologues of the human proteins were looked for. Primate proteomes are ranked according to the time of divergence between the primate and the human species, as provided in the TimeTree database, *Pan troglodytes* being on the left and *Protolemur simus* on the right.

As shown in Figure 3, while 89% of human proteins have homologues in all 27 proteomes of our reference database, 836 of them have homologues in all but one, 298 in all but two, *etc*, suggesting that the annotation of several reference proteomes is far from being complete.

**Figure 3:**
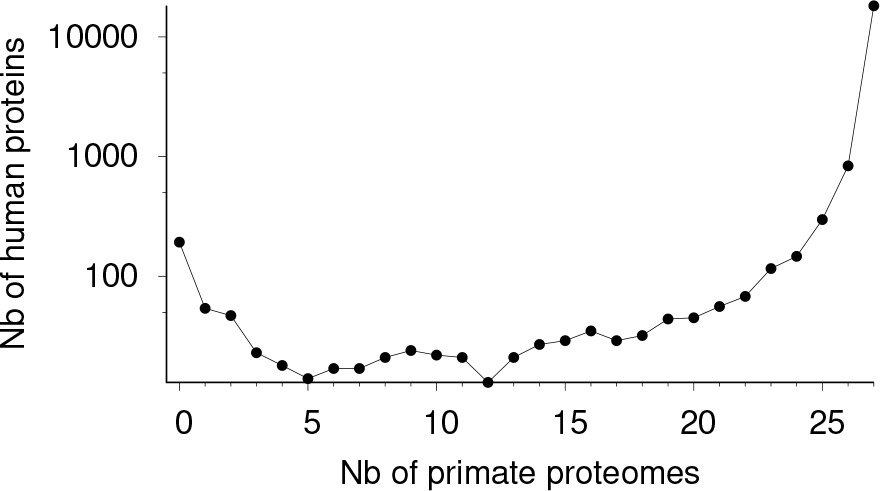
Number of human proteins with homologues found in a given number of primate proteomes. 193 human proteins have no homologue in the proteomes of the 27 primates considered.

Of course, if the annotation of the 27 proteomes considered were improved, or if more primate proteomes were added, the number of proteins found specific to the human species would keep on dropping. In order to partially take this expected trend into account, homologues of the 193 proteins found above were looked for in 52 Uniprot reference proteomes of other mammalian species, these other proteomes being chosen based on their high level of annotation, being all high-value outliers^5^, according to the Complete Proteome Detector [18].

Homologues were indeed found for 23 (12%) of them. However, for more than half of these cases (13/21) homologues were found in a single mammalian species *only*, as if the annotation of these proteins were intrinsically difficult. A possible reason is that these proteins are often short, with an average length of 106 ± 54 amino-acid residues.

### How many well-known ones ?

In Uniprot, the degree of knowledge, that is, the type of evidence that supports the existence of a protein is quantified through a number ranging between one (known at the protein level) and five (uncertain).

Among the 170 human-specific proteins identified above, only two are known at the protein level (see Table 1), according to Uniprot, namely, PACMP, the Poly-ADP-ribosylation-amplifying and CtIP-maintaining micropeptide [31], and SDIM1, the Stress-responsive DNAJB4-interacting membrane protein 1 [32]. Such a result is in sharp contrast with the fact that 90% of the human proteome is nowadays known at this level [33]. Note that PACMP is short (44 residues) and, as such, could have escaped annotation in the proteomes considered above. As a matter of fact, PACMP was included in Uniprot quite recently^6^.

**Table 1:** The 25 human specific proteins known either at the protein or transcript level. Most of them come from an open reading frame of a long non coding RNA. Proteins predicted to be 100% globular are in bold. The only protein with a fair quality of 3D structure prediction is underlined.

| Uniprot Identifier | Protein length | RNA Identifier<sup>a</sup> | Starting basis | Degree of knowledge<sup>b</sup> | Globularity<sup>c</sup> (domains) | pLDDT |
| --- | --- | --- | --- | --- | --- | --- |
| PACMP | 44 | URS0002617ABC | 139 | Protein | 0 (0) | ND<sup>d</sup> |
| **SDIM1** | 146 | **No** | – | Protein | 100 (1) | 29 |
| **Q68DW6** | 58 | URS00008B55A1 | 409 | Transcript | 100 (1) | 61 |
| <u>**CATR1**</u> | 79 | **No** | – | Transcript | 100 (1) | 83 |
| CF195 | 127 | URS00025F035E | 356 | Transcript | 97 (1) | 29 |
| YV004 | 128 | **No** | – | Transcript | 83 (1) | 41 |
| A4AS1 | 129 | URS000233F457 | 1298 | Transcript | 0 (0) | 49 |
| **YS039** | 131 | URS0002546BBE | 2824 | Transcript | 100 (1) | 36 |
| **HCP5** | 132 | URS00025BFBD4 | 372 | Transcript | 100 (1) | 39 |
| YP023 | 132 | URS000259F674 | 8 | Transcript | 61 (2) | 39 |
| **CA220** | 134 | URS00025BEBB1 | 359 | Transcript | 100 (1) | 39 |
| **CL036** | 138 | URS00025D0BC1 | 219 | Transcript | 100 (1) | 49 |
| **FEAS1** | 138 | URS00009BBF20 | 179 | Transcript | 100 (1) | 40 |
| YI001 | 140 | URS00025BBEEA | 315 | Transcript | 0 (0) | 38 |
| **YP033** | 140 | URS0002573B74 | 1427 | Transcript | 100 (1) | 31 |
| **YK004** | 145 | **No** | – | Transcript | 100 (1) | 38 |
| YS045 | 151 | URS0002339A91 | 77 | Transcript | 93 (1) | 41 |
| YF010 | 163 | URS00025B1F38 | 1769 | Transcript | 60 (2) | 34 |
| **YB035** | 174 | URS00025A88CD | 314 | Transcript | 100 (1) | 28 |
| YV008 | 177 | URS000258F499 | 37 | Transcript | 42 (1) | 26 |
| IDAS1 | 188 | URS0000A912F7 | 1 | Transcript | 99 (1) | 36 |
| YT009 | 211 | URS0000E60B4D | 490 | Transcript | 30 (1) | 41 |
| CQ077 | 243 | URS00021231B2 | 527 | Transcript | 91 (1) | 25 |
| CK072 | 251 | URS00025C6C2F | 242 | Transcript | 87 (1) | 27 |
| YU004 | 302 | URS0000EBC2BD | 89 | Transcript | 23 (1) | 26 |

<sup>a</sup>In the RNACentral database [25].
<sup>b</sup>According to Uniprot.
<sup>c</sup>Percentage of the protein length predicted to be globular, according to IUPred [14].
<sup>d</sup>PACMP was included in Uniprot after the release of the fourth version of the AlphaFold database [29].

On the other hand, while 23 of them are known at the transcript level (Table 1), 23 others are just predicted. Strikingly, 142 of them (71%) are coined uncertain, being annotated as dubious CDS or gene predictions, possible pseudogenes, *etc*.

Among the 25 human-specific proteins known either at the protein or transcript level, 21 (84%) are considered as being uncharacterized, meaning that they do not have any known or putative function.

On the other hand, 21 of them are found encoded by an open reading frame of a long non-coding RNA (lncRNA). Note that the growth of the number of reported non-coding RNA genes has been rapid, suggesting that the human catalogue may, in this respect, prove incomplete [10].

### How many globular ones ?

From the above results, it is tempting to speculate that most human-specific genes do not code for proteins and may instead be involved, for instance, in the regulation of gene expression [34]. However, it has recently been shown that lncRNAs show similar coding potential and sequence constraints than evolutionary young protein coding sequences [35]. Actually, up to 3330 human lncRNAs were found bound to ribosomes with active translation elongation [36].

It is thus necessary to assess the coding potential of lncRNAs. A straightforward way to do it is to predict how globular encoded proteins are expected to be [37]. As shown in Table 1, according to IUPred [14], 12 human-specific proteins known at the transcript level and encoded by lncRNA (60% of them) are predicted to be at least 90% globular, with a single structural domain. Note that all four human-specific proteins that are *not* known to be encoded by lncRNA (SDIM1, CATR1, YV004 and YK004) are predicted to be at least 83% globular, also with a single structural domain. On the other hand, since nearly 30% of regions within the proteome are expected to be disordered [30], other human-specific lncRNA could also encode genuine proteins.

### What about their structure ?

No homolog was found in the Protein Databank [38] for the human-specific proteins identified above. However, thanks to artificial intelligence algorithms, major progresses have recently been witnessed in the field of protein structure prediction [15, 16]. Moreover, such predictions have been performed on a large scale. Furthermore, they are nowadays available in public databases [29].

As a matter of fact, as illustrated in Figure 4, the tridimensional structure of most human proteins has been predicted with a high level of confidence by AlphaFold2 [29], the average pLDDT being over 90 for nearly 40% of them, and over 70 for more than 70% of them. Note that the structures of only 14% of human proteins are predicted with a very low confidence ((pLDDT) below 50).

**Figure 4:**
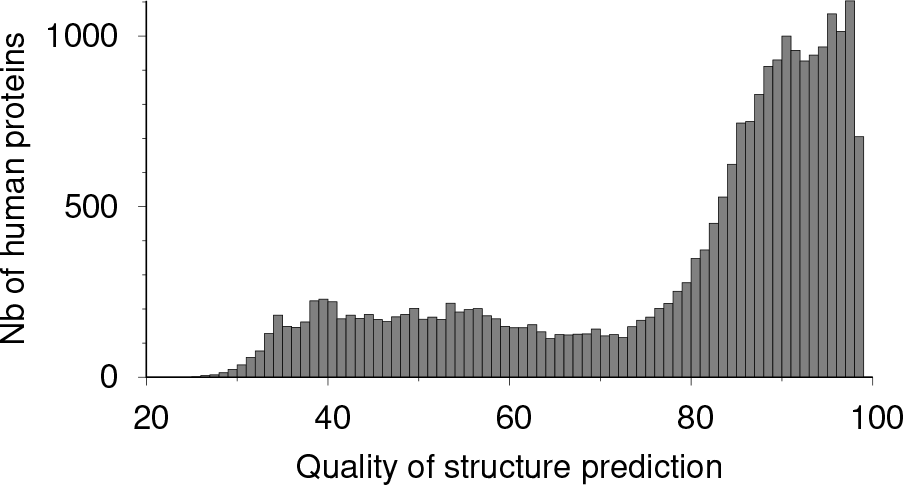
Number of human proteins with a given quality of 3D structure prediction, according to AlphaFold2. Quality ((pLDDT)) over 90 means: very high model confidence. Below 50: very low.

However, as shown in Table 1, among the 25 human-specific proteins known either at the protein or transcript level, AlphaFold2 is able to make a fair prediction ((pLDDT) = 83) in the case of a single of them only, namely, CATR1, the CATR tumorigenic conversion 1 protein [39, 40]. Note that its predicted structure is fairly simple, with a single, long α-helical segment (see Figure 5).

**Figure 5:**
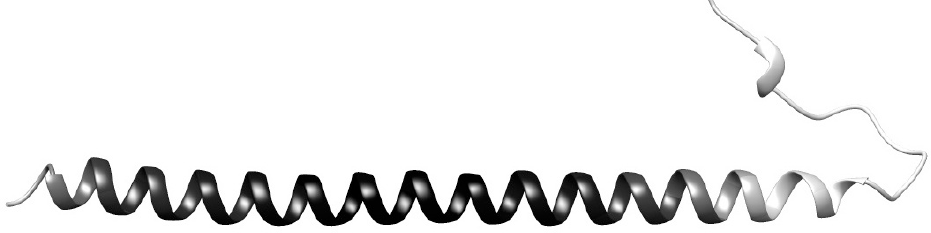
The structure of the human specific protein CATR1, as predicted by AlphaFold2 [15], with a fair level of confidence ((pLDDT)=83). The darker the higher the level of confidence. Drawn with Chimera [42].

Given that the structure of most proteins predicted to be globular by IUPred are poorly predicted by AlphaFold2 (Table 1), this result suggests that AlphaFold2 is of little help for predicting the structure of proteins with no known homolog [29], as for instance illustrated in a previous study of 362 eukaryotic proteomes [13].

## Conclusion

By looking for a lack of homologues in a reference database of 27 well-annotated proteomes of primates and 52 well-annotated proteomes of other mammals, 170 human-specific proteins were identified. However, most of them are coined dubious in Uniprot, casting doubts on 23 others that are just predicted. Indeed, given the efforts made in order to complete the annotation of the human proteome [10], it is less and less likely to have human proteins that are not known at the transcript level [33].

21 of the 25 human-specific proteins known either at the protein or transcript level are found encoded by an open reading frame of a lncRNA (Table 1). While a single of them, namely, PACMP, the Poly-ADP-ribosylation-amplifying and CtIP-maintaining micropeptide, is known at the protein level [31], twelve others are predicted to be at least 90% globular, with a single structural domain, suggesting that most, if not all, human-specific proteins are encoded by lncRNA.

As a matter of fact, some de novo genes have already been found to have a lncRNA origin [9, 35, 41]. Such lncRNA could come from intergenic open reading frames [37], which in turn may pop up randomly, becoming new functional proteins when they happen to confer selective advantages [41].

## Footnotes

1 On June 2023.

2 As found in Uniprot, on February 2023.

3 Standing for predicted local-distance difference test.

4 847 singletons.

5 On February 2023.

6 On October 2023, 12^*th*^.

## Notes

### Competing Interest Statement

The authors have declared no competing interest.

## References

[1] Toll-Riera, M., Bosch, N., Bellora, N., Castelo, R., Armengol, L., Estivill, X. & Alba, M.M. (2009). Origin of primate orphan genes: a comparative genomics approach. Mol. Biol. Evol. 26(3), 603–612.

[2] Cai, J.J. & Petrov, D.A. (2010). Relaxed purifying selection and possibly high rate of adaptation in primate lineage-specific genes. Gen. Biol. Evol. 2, 393–409.

[3] Ruiz-Orera, J., Hernandez-Rodriguez, J., Chiva, C., Sabidó, E., Kondova, I., Bontrop, R., Marqués-Bonet, T. & Albà, M.M. (2015). Origins of de novo genes in human and chimpanzee. PLoS Genet. 11(12), e1005721.

[4] Domazet-Loso, T. & Tautz, D. (2003). An evolutionary analysis of orphan genes in Drosophila. Genome Res. 13(10), 2213–2219.

[5] Wang, W., Yu, H. & Long, M. (2004). Duplication-degeneration as a mechanism of gene fission and the origin of new genes in Drosophila species. Nat. Genet. 36, 523–527.

[6] Lange, A., Patel, P.H., Heames, B., Damry, A.M., Saenger, T., Jackson, C.J., Findlay, G.D. & Bornberg-Bauer, E. (2021). Structural and functional characterization of a putative de novo gene in drosophila. Nat. Comm. 12(1), 1–13.

[7] Rich, A. & Carvunis, A.R. (2023). De novo gene increases brain size. Nature Ecology & Evolution 7(2), 180–181.

[8] Li, C.Y., Zhang, Y., Wang, Z., Zhang, Y., Cao, C., Zhang, P.W., Lu, S.J., Li, X.M., Yu, Q., Zheng, X. et al. (2010). A human-specific de novo protein-coding gene associated with human brain functions. PLoS Comput. Biol. 6(3), e1000734.

[9] An, N.A., Zhang, J., Mo, F., Luan, X., Tian, L., Shen, Q.S., Li, X., Li, C., Zhou, F., Zhang, B. et al. (2023). De novo genes with an lncRNA origin encode unique human brain developmental functionality. Nature Ecology & Evolution 7(2), 264–278.

[10] Amaral, P., Carbonell-Sala, S., De La Vega, F.M., Faial, T., Frankish, A., Gingeras, T., Guigo, R., Harrow, J.L., Hatzigeorgiou, A.G., Johnson, R. et al. (2023). The status of the human gene catalogue. Nature 622(7981), 41–47.

[11] Marques-Bonet, T., Ryder, O.A. & Eichler, E.E. (2009). Sequencing primate genomes: what have we learned? Annu. Rev. Genomics Hum. Genet. 10, 355–386.

[12] Juan, D., Santpere, G., Kelley, J.L., Cornejo, O.E. & Marques-Bonet, T. (2023). Current advances in primate genomics: novel approaches for understanding evolution and disease. Nat. Rev. Genet. 24(5), 314–331.

[13] Sanejouand, Y.H. (2023). On the unknown proteins of eukaryotic proteomes. J. Mol. Evol. 91, 492–501.

[14] Dosztányi, Z. (2018). Prediction of protein disorder based on IUPred. Protein Sci. 27(1), 331–340.

[15] Jumper, J., Evans, R., Pritzel, A., Green, T., Figurnov, M., Ronneberger, O., Tunyasuvunakool, K., Bates, R., Žídek, A., Potapenko, A. et al. (2021). Applying and improving AlphaFold at CASP14. Proteins: Struct., Funct., Bioinf. 89(12), 1711–1721.

[16] Jones, D.T. & Thornton, J.M. (2022). The impact of AlphaFold2 one year on. Nat. Methods 19(1), 15–20.

[17] UniProt Consortium (2017). Uniprot: the universal protein knowledgebase. Nucleic Acids Res. 45(D1), D158–D169.

[18] UniProt Consortium (2021). Uniprot: the universal protein knowledgebase in 2021. Nucleic Acids Res. 49(D1), D480–D489.

[19] Simão, F.A., Waterhouse, R.M., Ioannidis, P., Kriventseva, E.V. & Zdobnov, E.M. (2015). BUSCO: assessing genome assembly and annotation completeness with single-copy orthologs. Bioinformatics 31(19), 3210–3212.

[20] Kumar, S., Suleski, M., Craig, J.M., Kasprowicz, A.E., Sanderford, M., Li, M., Stecher, G. & Hedges, S.B. (2022). TimeTree 5: An expanded resource for species divergence times. Mol. Biol. Evol. 39(8), msac174.

[21] Altschul, S.F., Madden, T.L., Schäffer, A.A., Zhang, J., Zhang, Z., Miller, W. & Lipman, D.J. (1997). Gapped BLAST and PSI-BLAST: a new generation of protein database search programs. Nucleic Acids Res. 25, 3389–3402.

[22] Lobley, A., Swindells, M.B., Orengo, C.A. & Jones, D.T. (2007). Inferring function using patterns of native disorder in proteins. PLoS Comput. Biol. 3(8), e162.

[23] Lucas, S.J., Akpınar, B.A., Šimková, H., Kubaláková, M., Doležzel, J. & Budak, H. (2014). Next-generation sequencing of flow-sorted wheat chromosome 5D reveals lineage-specific translocations and widespread gene duplications. BMC genomics 15(1), 1–18.

[24] Schäffer, A.A., Aravind, L., Madden, T.L., Shavirin, S., Spouge, J.L., Wolf, Y.I., Koonin, E.V. & Altschul, S.F. (2001). Improving the accuracy of PSI-BLAST protein database searches with composition-based statistics and other refinements. Nucleic Acids Res. 29(14), 2994–3005.

[25] The RNAcentral Consortium (2015). RNA-central: an international database of ncRNA sequences. Nucleic Acids Res. 43, D123–D129.

[26] Dragomir, M.P., Manyam, G.C., Ott, L.F., Berland, L., Knutsen, E., Ivan, C., Lipovich, L., Broom, B.M. & Calin, G.A. (2020). FuncPEP: a database of functional peptides encoded by non-coding RNAs. Non-coding RNA 6(4), 41.

[27] Pajkos, M., Erdős, G. & Dosztányi, Z. (2023). The origin of discrepancies between predictions and annotations in intrinsically disordered proteins. Biomolecules 13(10), 1442.

[28] Necci, M., Piovesan, D. & Tosatto, S.C. (2021). Critical assessment of protein intrinsic disorder prediction. Nat. Methods 18(5), 472–481.

[29] Varadi, M., Anyango, S., Deshpande, M., Nair, S., Natassia, C., Yordanova, G., Yuan, D., Stroe, O., Wood, G., Laydon, A. et al. (2022). AlphaFold Protein Structure Database: Massively expanding the structural coverage of protein-sequence space with highaccuracy models. Nucl. Ac. Res. 50, D439–D444.

[30] Ruff, K.M. & Pappu, R.V. (2021). AlphaFold and implications for intrinsically disordered proteins. J. Mol. Biol. 433(20), 167208.

[31] Zhang, C., Zhou, B., Gu, F., Liu, H., Wu, H., Yao, F., Zheng, H., Fu, H., Chong, W., Cai, S. et al. (2022). Micropeptide PACMP inhibition elicits synthetic lethal effects by decreasing CtIP and poly(ADP-ribosyl)ation. Mol. Cell 82(7), 1297–1312.

[32] Lei, J.X., Cassone, C.G., Luebbert, C. & Liu, Q.Y. (2011). A novel neuron-enriched protein SDIM1 is down regulated in Alzheimer’s brains and attenuates cell death induced by DNAJB4 over-expression in neuro-progenitor cells. Molecular neurodegeneration 6(1), 1–16.

[33] Adhikari, S., Nice, E.C., Deutsch, E.W., Lane, L., Omenn, G.S., Pennington, S.R., Paik, Y.K., Overall, C.M., Corrales, F.J., Cristea, I.M. et al. (2020). A high-stringency blueprint of the human proteome. Nat. Comm. 11, 5301.

[34] Nahon, J.L. (2003). Birth of “human-specific” genes during primate evolution. Genetica 118, 193–208.

[35] Ruiz-Orera, J., Messeguer, X., Subirana, J.A. & Alba, M.M. (2014). Long non-coding RNAs as a source of new peptides. elife 3, e03523.

[36] Lu, S., Zhang, J., Lian, X., Sun, L., Meng, K., Chen, Y., Sun, Z., Yin, X., Li, Y., Zhao, J. et al. (2019). A hidden human proteome encoded by “non-coding” genes. Nucleic Acids Res. 47(15), 8111–8125.

[37] Papadopoulos, C., Callebaut, I., Gelly, J.C., Hatin, I., Namy, O., Renard, M., Lespinet, O. & Lopes, A. (2021). Intergenic ORFs as elementary structural modules of de novo gene birth and protein evolution. Genome Res. 31(12), 2303–2315.

[38] Kouranov, A., Xie, L., De La Cruz, J., Chen, L., Westbrook, J., Bourne, P.E. & Berman, H.M. (2006). The RCSB PDB information portal for structural genomics. Nucl. Ac. Res. 34, D302–D305.

[39] Li, D., Noyes, I., Shuler, C. & Milo, G.E. (1995). Cloning and sequencing of CATR1. 3, a human gene associated with tumorigenic conversion. Proc. Natl. Acad. Sci. USA 92(14), 6409–6413.

[40] Li, D., Sun, X.L., Casto, B., Fang, J., Theil, K., Glaser, R. & Milo, G. (1998). Epstein–Barr virus growth-transformed cells are converted to malignancy following transfection of a 1.3-kb CATR1 antisense construct independent of a change in the level of c-myc expression followed by a 8;14 chromosomal translocation. Proc. Natl. Acad. Sci. USA 95(9), 4894–4899.

[41] Ruiz-Orera, J., Villanueva-Cañas, J.L. & Albà, M.M. (2020). Evolution of new proteins from translated sORFs in long non-coding RNAs. Exp. Cell Res. 391(1), 111940.

[42] Pettersen, E.F., Goddard, T.D., Huang, C.C., Couch, G.S., Greenblatt, D.M., Meng, E.C. & Ferrin, T.E. (2004). UCSF chimera: a visualization system for exploratory research and analysis. J. Comput. Chem. 25(13), 1605–1612.

